# Predictive orientation remapping maintains a stable retinal percept

**DOI:** 10.1101/193250

**Authors:** T. Scott Murdison, Gunnar Blohm, Frank Bremmer

**Author notes:** **Competing interests:** The authors declare no competing financial interests. **Abbreviated title:** Retinal predictive orientation remapping. **Contact:** Gunnar Blohm, Centre for Neuroscience Studies (CNS), Queen’s University, Botterell Hall, room 229,18 Stuart St., Kingston, Ontario, Canada, K7L3N6.

## Abstract

Despite motion on the retina with every saccade, we perceive the world as stable. But whether this stability is a result of neurons constructing a spatial map or continually remapping a retinal representation is unclear. Previous work has focused on the perceptual consequences of shifts in the horizontal and vertical dimensions, but torsion is another key component in ocular orienting that – unlike horizontal and vertical movements – produces a natural misalignment between spatial and retinal coordinates. Here we took advantage of oblique eye orientation-induced retinal torsion to examine perisaccadic orientation perception. We found that orientation perception was largely predicted by the retinal image throughout each trial. Surprisingly however, we observed a significant presaccadic remapping of the percept consistent with maintaining a stable (but spatially inaccurate) retinotopic perception throughout the saccade. These findings strongly suggest that our seamless perceptual stability relies on retinotopic signals that are remapped with each saccade.

## Introduction

We move our eyes all the time, and with every movement we induce massive shifts of the retinal projection. Despite this motion, we can keep track of both the locations and features (e.g. orientation) of objects in space. To achieve such stability, the perceptual system is thought to compensate for each eye movement using predictive remapping. Separate recordings from distinct retinotopic areas have revealed that receptive fields (RFs) presaccadically modulate their spatial tuning by either shifting (Zirnsak, Steinmetz, Noudoost, Xu, & Moore, 2014) or expanding (J. Duhamel, Colby, & Goldberg, 1992; Wang et al., 2016) towards the target. Consequently, these presaccadic RF modulations are assumed to be involved in the maintenance of perceptual stability, though how is unclear. Two potential explanations that have garnered some recent debate (D. Burr, Tozzi, & Morrone, 2007; J. R. Duhamel, Bremmer, Ben Hamed, & Graf, 1997; Harrison & Bex, 2014; Harrison, Mattingley, & Remington, 2012; Melcher, 2005; Morris, Bremmer, & Krekelberg, 2016; Turi & Burr, 2012; Zimmermann, Burr, & Morrone, 2011; Zimmermann, Morrone, Fink, & Burr, 2013; Zirnsak & Moore, 2014) are that either these RF modulations predictively remap a retinotopic representation purely in compensation for the upcoming retinal motion or they are involved in constructing a stable spatial map of the visual scene.

Previous remapping work has only considered two-dimensional (2D) motion on the retina when in fact, shifts in the third, torsional dimension (i.e., around a rotation axis parallel to the line of sight) is also present during almost any eye movement and is a key component of ocular orienting. For example, retinal torsion can be induced by ocular counter-roll during head roll (Blohm & Lefèvre, 2010; Murdison, Paré-Bingley, & Blohm, 2013), by the natural tilt of Listing’s plane (Blohm, Khan, Ren, Schreiber, & Crawford, 2008), or by simply manipulating the geometry of the retinal projection using oblique gaze orientations (Blohm & Lefèvre, 2010) (which, importantly, does not require any mechanical torsion of the eyeball). Historically, differentiating between the retinal and spatial models has been impossible without linking remapping to exogenous factors such as visual attention (Harrison et al., 2012; Mathôt & Theeuwes, 2010; Rolfs, Jonikaitis, Deubel, & Cavanagh, 2011), the motion (Turi & Burr, 2012) or tilt (Melcher, 2007) after-effects, or object features (Julie D. Golomb, L’Heureux, & Kanwisher, 2014; Harrison & Bex, 2014). Conveniently, torsion provides a natural misalignment between retinal and spatial coordinates for which the perceptual system must directly compensate. Here, we geometrically induced torsional shifts by projecting a frontoparallel stimulus onto the retina during movements to and from oblique eye orientations (oblique orientation-induced retinal torsion, ORT, Fig. 1A). Past work has found that ORT influences orientation perception in a retinally predicted way during fixation (Haustein & Mittelstaedt, 1990; Nakayama & Balliet, 1977), yet no study has examined how ORT affects orientation perception during ongoing eye movements.

There are three possible perceptual outcomes of predictive remapping across torsional shifts. First, there might be no predictive remapping, with orientation perception adhering to ORT (Haustein & Mittelstaedt, 1990; Nakayama & Balliet, 1977) throughout the movement (null model). Second, the perceptual system might use an estimate of the future retino-spatial geometry to presaccadically and predictively tilt perception *towards* the final ORT, ahead of the eyes (retino-spatial model, Fig. 1B). Third, the perceptual system might presaccadically tilt perception *away* from the final orientation, allowing a retinotopic perception to move with the eyes (purely retinal model, Fig. 1C). Here we provide strong evidence in support of the purely retinal model using ORT during a perisaccadic orientation perception task.

**Figure 1.**
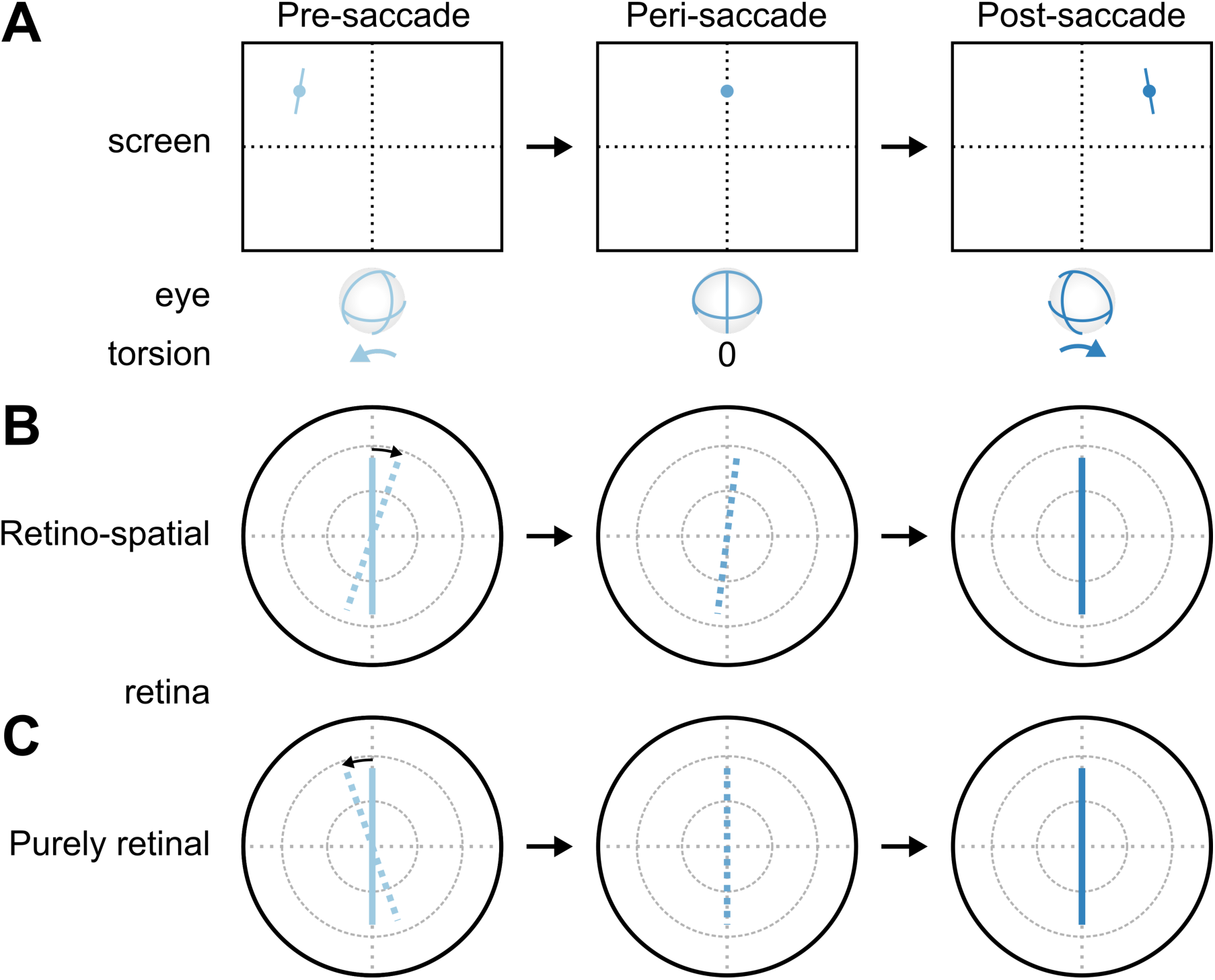
Geometry of predictive remapping models. **A** Example ORT during oblique-to-oblique horizontal saccades. Note that ORT magnitude is exaggerated for illustration purposes. **B** Retinotopic representation of the retino-spatial predictive model. Solid lines represent the actual retinal projection of the stimulus while dotted lines represent the corresponding percept. **C** Retinotopic representation of the purely retinal predictive model.

## Results and discussion

We directly investigated how ORT influences orientation perception across saccades using a novel retinal feature remapping paradigm in complete darkness. Participants performed either the test version of the task between oblique gaze locations (inducing ORT) or the control version of the task along the horizontal screen meridian (Fig. 2A). They began each trial by fixating a target on the left side of the screen (Fig. 2B). 300 ms later, a second target was illuminated on the opposite (right) side of the screen. After a randomized fixation duration, we briefly presented an oriented bar stimulus in one of seven different orientations rotated from vertical at the current gaze location. At the end of the trial, participants reported their perception of the stimulus orientation relative to gravity (clockwise, CW, or counter-clockwise, CCW). This paradigm allowed us to reliably compute each participant’s psychometric function with a fine time resolution throughout the saccade. A fixation version of the task in which we presented the same stimulus at one of six possible fixation locations (three along each test or control trajectory – left, center and right) allowed us to account for any perceptual effects during stable fixation. We also measured each participant’s natural Listing’s plane to account for any natural ocular torsion when making predictions using our retinal model (Table 1 in Methods).

**Figure 2.**
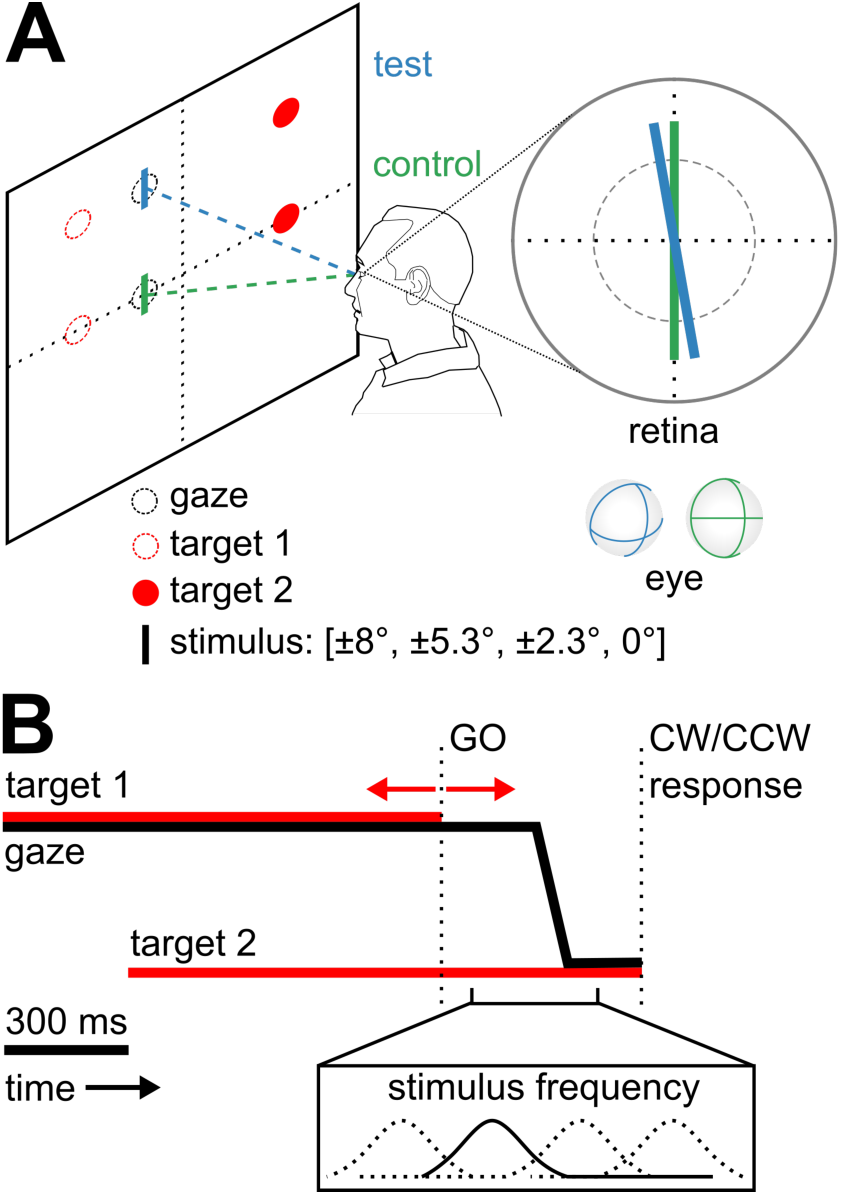
Paradigm and task timing. **A** Illustration demonstrating rotational effects induced on retina due to oblique eye orientations while participants do task in either the test or control condition. Note that these retinal rotations are exaggerated for illustration purposes. **B** Schematic showing task timing and stimulus presentation frequency distributions. Bidirectional arrows represent 200 ms time window within which we randomly varied “go” cue.

**Table 1.**
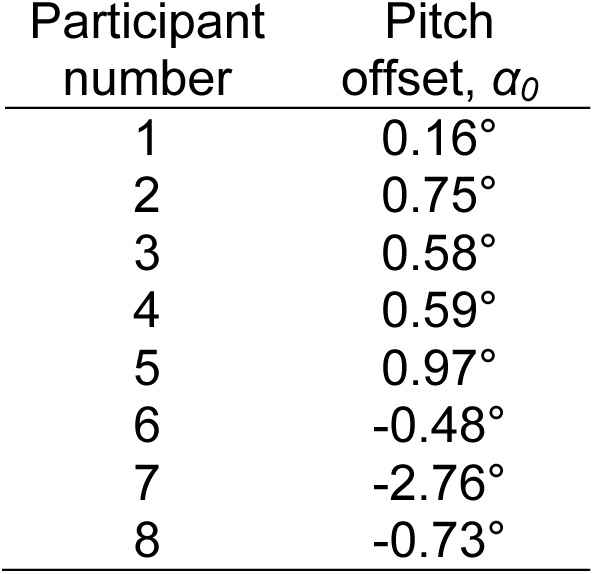
Identified Listing’s plane tilt for each participant.

| Participant<br>number | Pitch<br>offset, $\alpha_0$ |
| --- | --- |
| 1 | $0.16^\circ$ |
| 2 | $0.75^\circ$ |
| 3 | $0.58^\circ$ |
| 4 | $0.59^\circ$ |
| 5 | $0.97^\circ$ |
| 6 | $-0.48^\circ$ |
| 7 | $-2.76^\circ$ |
| 8 | $-0.73^\circ$ |

We examined the performance of each participant as a function of trial time (aligned to saccade onset) revealing orientation perception throughout any given trial, and we compared perceptions to the prediction of a retinal model. Participants had clear perceptual differences (Fig. 3) between the start (light shades) and end (dark shades) of the saccade, but these differences were most pronounced for test trials. As the eyes moved across screen we found that the perceptual changes were captured by the retinal model predictions during test trials (pooled regression analysis across 4° on-screen bins, *n* = 12, *slope* = 0.87, *R*^2^ = 0.7, *p* < 0.01), and matched perceptions during fixation at the extreme time points (paired T-test for points of subjective equality, *t*(15) = 1.52, *p* = 0.15, and for just-noticeable differences, *t*(15) = 0.47, *p =* 0.65), indicating that they behaved consistently during periods when the eyes were stationary, regardless of the behavioral context (fixation vs. saccade trials).

**Figure 3.**
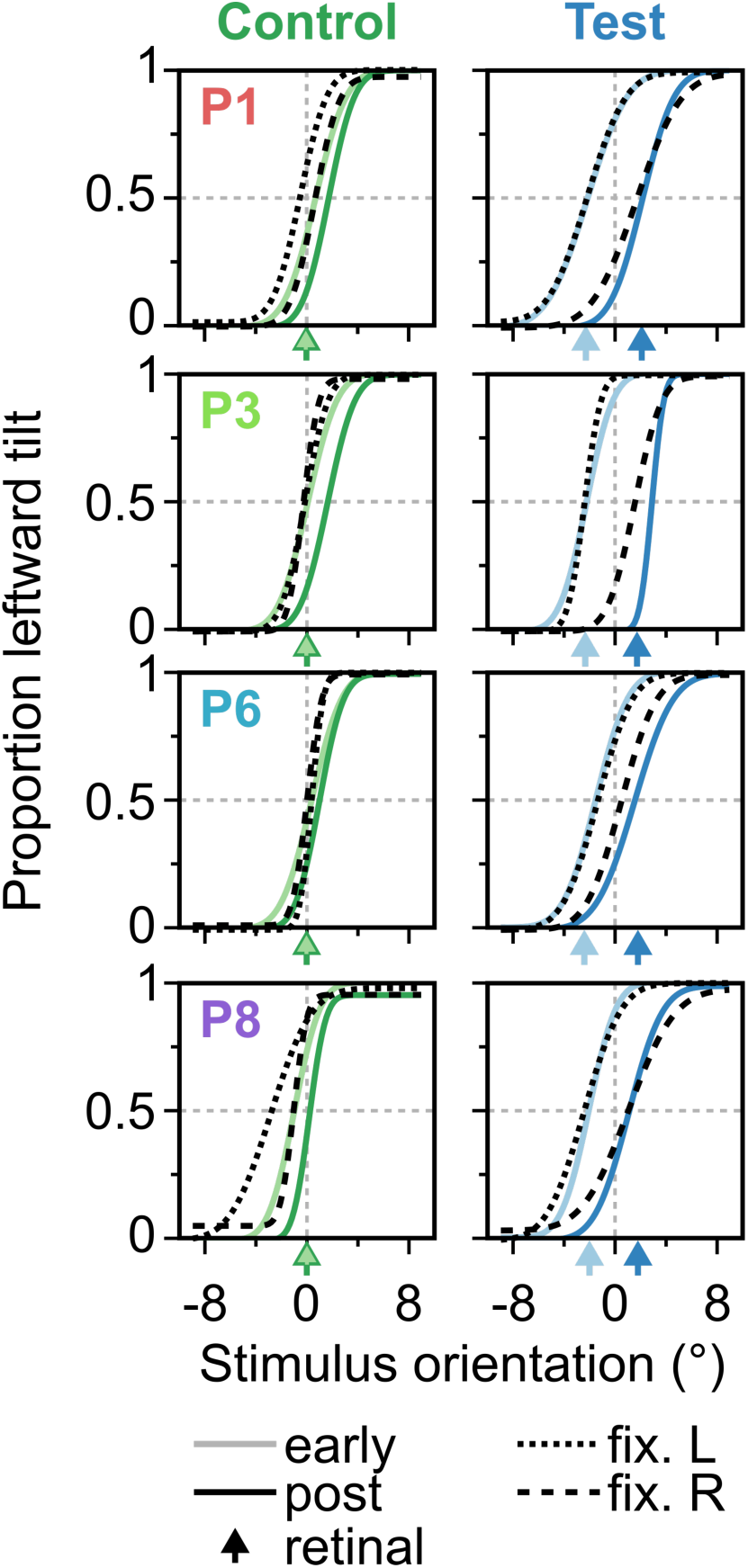
Psychometric curves for four sample participants. Early (trial start until 50 ms presaccade; light shades) and late (100 ms postsaccade until trial end; dark shades) psychometric function fits are shown alongside the fixation experiment results for left (dotted) and right (dashed) targets. Color-matched arrows represent retinal predictions for 50% thresholds during each time bin.

After pooling the data across participants we were able to attain a time resolution of 15 ms for which we could compute the bin-wise psychometric functions, extracting the points of subjective equality (PSEs) to quantify the psychophysical biases and the just-noticeable differences (JNDs) to quantify the corresponding precision. These time-resolved biases (PSEs with 95% confidence intervals) are shown alongside the retinal predictions (dashed lines) for both control (green) and test trials (blue) relative to saccade onset (Fig. 4A). Psychophysical biases depended on whether participants performed control or test trials. Throughout control trials, perceptual biases followed the retinally predicted perception, with excursions from the retinal prediction occurring upon, but not prior to, saccade onset. Throughout test trials however, orientation perception was biased towards the retinal prediction throughout the movement, with the exception of a significant perceptual rotation immediately prior to the movement onset. Using the pooled data, the effect began approximately 50 ms prior to the movement (grey shaded window; inset), consistent with the timing of both attentional (Harrison et al., 2012; Rolfs et al., 2011) and RF shifts observed in retinotopic areas (Wang et al., 2016; Zirnsak et al., 2014). Furthermore, this deviation went in the direction opposite to the upcoming shift in ORT in a manner consistent with maintaining the retinotopic orientation throughout the upcoming movement, matching the purely retinal model.

**Figure 3.**
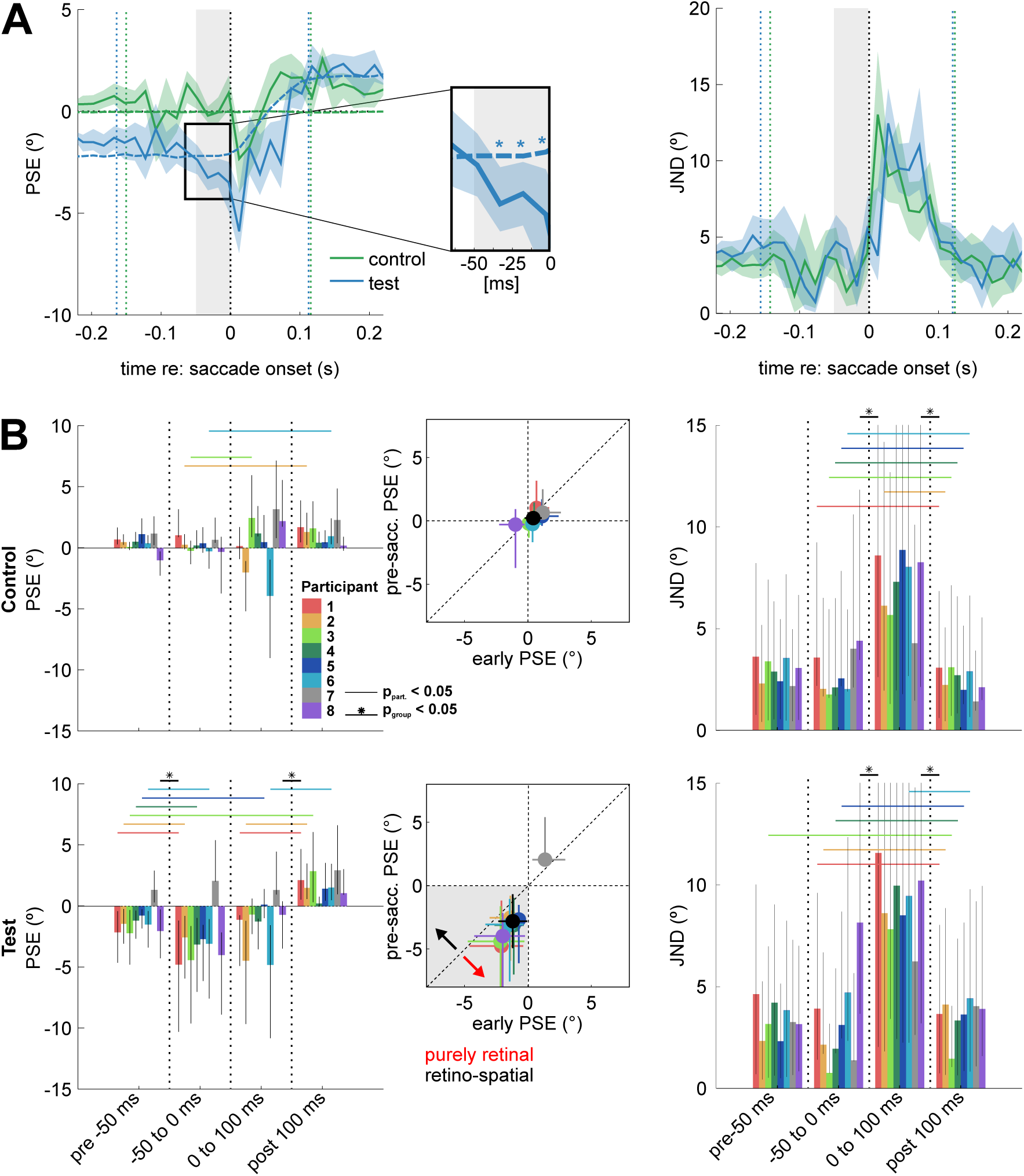
Pooled and participant-level biases. **A** Pooled PSEs (*left column*) and JNDs (*right column*) for control (green) and test trials (blue), plotted alongside the retinal predictions (color-matched dashes) over time. Inset reveals significant presaccadic perceptual rotation for test PSEs (asterisks). **B** Participant-level PSEs and JNDs, binned into early (*t* < -50 ms), presaccadic (-50 ms < *t* < 0 ms), perisaccadic (0 ms < *t* < 100 ms) and post-saccadic time bins (*t* > 100 ms), aligned to saccade onset for control (*top row*) and test trials (*bottom row*). Participant-level significant effects are shown by color-matched bars crossing bin thresholds (vertical dotted lines), and group-level significant effects are shown by bold black crossing lines and black asterisks. Insets (*center column*) reveal direct comparisons between PSEs in presaccadic (ordinate) and early time bins (abscissa). Within the test inset, shaded quadrant represents the retinal hypothesis for either time epoch, and arrows represent direction of retino-spatial (black) or purely retinal (red) remapping. Black circles and error bars represent across-participant means and standard deviations.

We next determined if this observed effect during test trials was simply a phenomenological effect of pooling the data across participants (Fig. 4B). We separated each participant’s data into four separate time bins representing characteristic time epochs during any given trial: 1. early fixation (trial start to 50 ms prior to onset); 2. Presaccadic (50 ms prior to saccade onset); 3. Perisaccadic (saccade onset to 100 ms later); and 4. Postsaccadic (100 ms post saccade onset until trial end). Using these binned data, we observed the same presaccadic bias shift on the group level for test trials (paired t-test, *t*(7) = -4.33, *p* < 0.01), indicating that it was not due to pooling data across participants. We varied the presaccadic bin size as much as participants’ time resolutions allowed, and found qualitatively identical group-level presaccadic remapping effects up to 40 ms prior to onset (not shown here). Finally, as these bias shifts could potentially be simply explained by a less precise perception, we also examined the time-resolved changes in precision. We did this with JNDs in an identical way (Fig. 4A and B, right column), and found that they only increased perisaccadically (paired t-tests, all transsaccadic *p* < 0.01), as expected from retinal blurring and/or saccadic suppression (Bremmer, Kubischik, Hoffmann, & Krekelberg, 2009; D. C. Burr, Morrone, & Ross, 1994), but presaccadic precision was not different from precision during fixation. Thus, presaccadic perceptual shifts could not be explained by a decrease in perceptual precision.

We found that ORT, which is not corrected for during fixation (Haustein & Mittelstaedt, 1990; Nakayama & Balliet, 1977), is predictively remapped across saccades in an orientation perception task. Instead of updating the perception ahead of the eye movement using an estimate of the spatial geometry at the final gaze location (retino-spatial model), the presaccadic shifts we observe instead are compensatory for the future ORT, allowing the retinotopic orientation to be maintained while the eyes move (purely retinal model). This key finding is in agreement with recent psychophysical work (Julie D. Golomb et al., 2014; Rolfs et al., 2011).

The predictive orientation shifts we observed are also consistent with the hypothesis that presaccadic RF shifts in retinotopic areas contribute to the stability of visual perception (Wang et al., 2016; Zirnsak et al., 2014). Consequently, elucidating the neural substrate of these perceptual shifts could potentially reconcile contrasting shifting (Zirnsak et al., 2014) and expansion (Wang et al., 2016) RF models of predictive remapping. Our psychophysical results predict that the activity of orientation-selective retinotopic neurons involved in predictive remapping should also exhibit torsion-induced modulations. Such predictive orientation-selective modulations might, for example, be seen in extrastriate visual cortex (Nakamura & Colby, 2002), lateral intraparietal area (J. Duhamel et al., 1992; Wang et al., 2016) or frontal eye fields (Zirnsak et al., 2014), perhaps from retinotopic corollary discharge signals arising in superior colliculus (Sommer & Wurtz, 2004) projected via the mediodorsal thalamus (Sommer & Wurtz, 2004; Zimmermann & Bremmer, 2016).

The implication that the brain expends computational energy with each eye movement to predictively remap a (spatially incorrect) retinal perception is seemingly paradoxical; after all, in theory the brain has access to all the self-motion signals required to compensate for retinal blurring and/or retino-spatial misalignments. However, compensating for self-motion requires either updating of a non-spatial (e.g. retinal) representation (D. Y. P. Henriques, Klier, Smith, Lowy, & Crawford, 1998; Medendorp, Van Asselt, & Gielen, 1999; Murdison et al., 2013) or subjecting sensory signals to reference frame transformations (Blohm & Crawford, 2007; Blohm & Lefèvre, 2010; Murdison, Leclercq, Lefèvre, & Blohm, 2015) to achieve spatial accuracy. As both updating (Medendorp et al., 1999) and reference frame transformations appear to be stochastic (Alikhanian, Carvalho, & Blohm, 2015; Jessica K Burns, Nashed, & Blohm, 2011; Jessica Katherine Burns & Blohm, 2010; Schlicht & Schrater, 2007; Sober & Sabes, 2003) processes, retinotopic signals might provide high acuity sensory information on which to base working memory (Julie D Golomb, Chun, & Mazer, 2008), perception (Jessica K Burns et al., 2011; Rolfs et al., 2011) and movement generation (Schlicht & Schrater, 2007; Sober & Sabes, 2003) explicitly requiring a reference frame transformation.

The apparent dominance of retinotopic signals we observed during saccades is consistent with a growing body of psychophysical (Julie D. Golomb et al., 2014; Murdison et al., 2013; Rolfs et al., 2011; Zirnsak, Gerhards, Kiani, Lappe, & Hamker, 2011) and electrophysiological (Colby, Duhamel, & Goldberg, 1995; J. Duhamel et al., 1992; J. R. Duhamel et al., 1997; Wang et al., 2016; Zirnsak et al., 2014) evidence. Indeed, participants are better at recalling the retinotopic locations of stimuli across saccades compared to their spatial locations, which are degraded with each subsequent eye movement (J. D. Golomb & Kanwisher, 2012). Additionally, attention appears to be allocated in retinotopic coordinates (J. D. Golomb, Nguyen-Phuc, Mazer, McCarthy, & Chun, 2010; Julie D Golomb et al., 2008; Yao, Ketkar, Treue, & Krishna, 2016) and there is evidence that its locus shifts to the retinotopic target of upcoming saccades (Mathôt & Theeuwes, 2010; Rolfs et al., 2011). Memorized targets for movement also appear to be encoded retinotopically, as observed during saccades (Inaba & Kawano, 2014), smooth pursuit (Murdison et al., 2013) and reaching (Batista, Buneo, Snyder, & Andersen, 1999; D. Y. Henriques, Klier, Smith, Lowy, & Crawford, 1998; Medendorp et al., 1999). Together with this past work, our findings indicate that reliable retinal signals are paramount to maintaining a stable world percept during self-motion. Corollary to this claim is that the natural statistics of the visual environment appear to play a more central role than extraretinal signals in forming that world percept on the millisecond timescale. As such, investigations into the temporal stability of stimuli and retinal scene characteristics required during saccades for a spatially correct perception are logical extensions of this work.

For the first time, we have shown the orientation-specific perceptual consequences of shifts in the torsional dimension during saccades. Together with previous work (Wang et al., 2016; Zirnsak & Moore, 2014; Zirnsak et al., 2014), our current findings imply that the perceptual system faithfully maintains a retinotopic representation by predictively remapping across both translational and torsional retinal shifts. In the midst of motion on the retina with each exploratory eye movement, it appears that this predictive remapping underlies the seamless stability that is a hallmark of our perceptual experience.

## Methods

### Participants

Eight adults with normal or corrected to normal vision performed the experiment (3 female, age range 20-30 years). Participants were paid for their participation and were all naïve to the purpose of the experiment, and all had previous experience with psychophysical experiments involving video eye tracking. Each participant gave informed written consent prior to the experiment. All procedures used in this study conformed to the Declaration of Helsinki.

### Materials

Stimuli were computer-generated using the Psychophysics Toolbox (Brainard, 1997) within Matlab (The Mathworks, Inc., Natick, Massachusetts), and were projected onto a large 120 cm (81°) x 90 cm (65.5°) flat screen by means of a DS+6K-M Christie projector (Christie Digital, Cypress, California) at a frame rate of 120 Hz and a resolution of 1152 x 864 pixels. Participants sat in complete darkness 70 cm away from the screen, and a table-mounted chin rest supported their heads. The complete darkness was required to prevent subjects perceive a compression of space, which might have confounded our data (Krekelberg, Kubischik, Hoffmann, & Bremmer, 2003; Lappe, Awater, & Krekelberg, 2000; Morrone, Ross, & Burr, 1997). Eye movements were recorded using an infrared video-based Eyelink II (SR Research, Ottawa, Ontario) that was attached to the chin rest, providing a table-fixed head strap that kept each participant’s head in a constant position throughout each experimental session. The screen was viewed binocularly and eye position was sampled at 500 Hz. Prior to each block, participants performed a 13-point calibration sequence over a maximum eccentricity of 25°. The eye to which the perceptual stimulus was fovea-locked for each block was selected based on calibration performance. Drift correction was performed offline every 10 trials, based on a central fixation position. To ensure precise temporal measurement of trial start and stimulus presentation, we positioned a photosensitive diode over the lower left corner of the screen, where we flashed a white patch of pixels both at the start of each trial and at the presentation of the oriented bar stimulus. This part of the experimental apparatus was occluded from the view of the participant. After calibration for constant data acquisition delays, the photosensitive diode’s voltage spikes provided reliable estimates of each trial’s time-course (within a precision of approximately 2 ms).

### Procedure

Participants performed a two-alternative, forced choice (2AFC) perceptual task in which they made large horizontal saccades between targets 40° apart either along a 20° vertically eccentric horizontal axis (test trials) or along the horizontal meridian of the screen (control trials; Fig. 2A). Importantly, test trials induced ORT throughout the eye movement. Participants began each trial by fixating the initial 0.3° diameter dot on the left side of the screen (at -20°), and indicated with a key press that they were prepared to start the trial (Fig. 2B). 300 ms later, a 0.3° diameter target was illuminated 40° to the right on the opposite side of the screen (at +20°). After a randomly selected duration (400-600 ms), the initial target was extinguished, representing the participant’s “go” cue. At some point in time, either immediately before saccade onset (~250 ms prior), during the saccade (average saccade duration ~120 ms) or after the saccade, we presented an oriented bar stimulus in one of 7 different orientations (from -8° to +8° rotated from vertical). For each trial, the exact time at which we presented the stimulus was chosen randomly from one of four 200 ms-width Gaussians, linearly spaced from the average reaction time (based on a 10-trial moving window) to 100 ms after, approximating the end of the movement. After the participant’s eyes had landed on the saccade target, they were asked to respond with a key press representing their perception of the stimulus orientation (counter-clockwise or clockwise perceptions). The trial ended after participants made their selection. This paradigm allowed us to reliably compute each participant’s psychometric function with a fine time resolution throughout a saccade.

Participants also performed a fixation version of the same task in which they fixated one of six randomly selected locations (-20°, 0° or +20° horizontal along either the 0° or 20° screen meridian) and we flashed the identical stimulus at the fixation location for a single frame (8.3 ms). After the stimulus flash participants responded with a key press indicating their perception of its orientation, identically to the first experiment.

### Identifying Listing’s plane for each participant

Finally, to correctly compute the retinal model predictions we measured each participant’s individual Listing’s plane (Table 1) using photographs taken during fixation at each of 10 orientations on the screen (rectangular grid in the upper half of the screen along 0° and 20° meridians, with five equally-spaced orientations along each horizontal and 20° eccentricity). From these photographs we extracted the natural ocular torsion based on the irises compared between the central orientation (0°, 0°) and eccentric locations, using an algorithm developed by Otero-Millan and colleagues (Otero-Millan, Roberts, Lasker, & Zee, 2015) modified for still images and implementation in Matlab.

### Analysis

All analyses were performed using custom Matlab code (The Mathworks, Natick, Massachusetts) and psychometric functions were fit using the Psignifit toolbox (Wichmann & Hill, 2001). Each participant performed 2080 trials in total, following a Gaussian distribution of presented stimulus orientations. Each performed a minimum of 221 repetitions for each of the most extreme bar orientations (+/-8°); conversely, for 0° bar orientations, they performed a maximum of 369 repetitions. These repetitions allowed us to be confident in our psychometric fits while not extending the sessions by oversampling easy trials. Trials containing blinks, loss of eye tracking, no saccades, hypometric or inaccurate saccades (<25° amplitude or beyond 10° radius from target), or with reaction times greater than 1.5 s were all removed from the dataset (20% of all trials). Group-level statistics were computed using paired Student t-tests, and participant-level and pooled analyses were performed using the bootstrapped 95% confidence intervals determined from Monte Carlo simulations during the psychometric curve fitting.

## Acknowledgments

The authors want to thank Dr. Dominic Standage for his helpful comments on the manuscript, as well as the participants for their kind participation. This work was supported by DFG (IRTG/CREATE-1901-The Brain in Action, Germany), NSERC (Canada), CFI (Canada), the Botterell Fund (Queen’s University, Kingston, ON, Canada) and ORF (Canada). TSM was also supported by DAAD (Germany).

